# Mice detect and discriminate temporal patterns of mesoscale optogenetic cortical stimulation

**DOI:** 10.64898/2026.09.17.752395

**Authors:** Gabrielle Stephens, Nikolas A. Francis

## Abstract

Optogenetic stimulation is widely used to establish causal links between neural activity, behavior, and sensory perception, yet animals may also perceive the neural stimulation itself, a phenomenon termed “optoception.” We hypothesized that cortical optogenetic stimulation would generate an artificial percept that could be distinguished from externally evoked sensations and characterized psychometrically in an operant task. We tested this hypothesis using mesoscale cortical photostimulation delivered by a 3 mm, 459 nm LED implanted over auditory cortex. Four mice expressing Channelrhodopsin-2 (ChR2) in excitatory neurons learned a head-fixed go/no-go task in which 20 Hz cortical photostimulation served as the target. ChR2-positive mice detected cortical photostimulation relative to LED-off catch trials, distinguished it from both an external LED matched in wavelength and pulse rate and from an acoustic noise matched in amplitude-modulation rate, and discriminated between 20- and 10-Hz cortical photostimulation. Reducing implanted-LED optical power initially lowered rate-discrimination sensitivity, which then increased across training sessions, demonstrating perceptual learning. In two mice tested with 11 randomly interleaved rates spanning 10-20 Hz, lick probability followed a saturating rate-dependent function. Two ChR2-negative littermates failed to acquire photostimulation-locked behavioral responding. Together, these findings show that mesoscale cortical optogenetic stimulation generates a graded, learnable artificial percept whose behavioral readout depends on photostimulation rate and optical power. This single-LED preparation provides a cost-efficient and wavelength-extensible platform for psychometric analysis of optoception.

**SIGNIFICANCE STATEMENT:** Optogenetic stimulation can alter sensory processing, but the neural perturbation itself may also become a perceptual cue. We show that mice detect mesoscale cortical photostimulation delivered by a 3 mm implanted LED, distinguish it from matched external light and sound, and discriminate differences in photostimulation rate. When implanted-LED optical power was reduced, discrimination sensitivity initially declined and then increased across training sessions, revealing perceptual learning of optoception. Thus, mesoscale cortical photostimulation generates a graded internal signal whose behavioral readout depends on stimulation rate, optical power, and experience. A single implanted LED provides a cost- efficient and wavelength-extensible assay for measuring optogenetic perception.

## INTRODUCTION

Optogenetic stimulation permits millisecond-scale control of genetically defined neurons and has become a central method for testing causal links between neural activity and sensory perception (Boyden et al., 2005; Carrillo-Reid et al., 2019; Deisseroth, 2015; Fenno et al., 2011; Huber et al., 2008; Kang et al., 2025; Kim et al., 2017; Marshel et al., 2019). However, perceptual changes during optogenetic stimulation may arise because the animal detects the artificial stimulation itself and learns that it predicts a behavioral outcome, rather than solely because the manipulation alters natural sensory processing. Luis-Islas et al. (2022) termed this perception of optogenetic brain perturbations “optoception.” Behaviorally reportable optogenetic stimulation has been demonstrated at several spatial scales and in multiple sensory systems. Mice can detect activation of sparse populations of layer 2/3 neurons in barrel cortex (Huber et al., 2008), and training can reduce the number of neurons required for detection (Dalgleish et al., 2020). Mice can also discriminate the number of photostimulation pulses delivered to somatosensory cortex (Pancholi et al., 2023). Electrical microstimulation studies further showed that imposed cortical timing can support quantitative psychophysical judgments (Romo et al., 2000; Romo et al., 1998; Yang et al., 2008). Patterned optogenetic activation of visual or olfactory ensembles can bias or generate learned perceptual reports (Carrillo-Reid et al., 2019; Chong et al., 2020; Gill et al., 2020; Histed & Maunsell, 2014; Marshel et al., 2019), and patterned transcranial stimulation can generate artificial percepts (Wu et al., 2026). Together, these findings suggest that optoception preserves information about the intensity, spatial organization, and temporal pattern of a neural perturbation and can exhibit experience-dependent plasticity.

Auditory cortex provides a useful system for testing whether temporally structured optogenetic stimulation is directly accessible to behavior. Auditory cortical areas form interconnected processing networks (Hackett, 2011) and encode temporal modulation in sound (Putnam et al., 2026; Yin et al., 2011). Activity in auditory cortex is modulated during sound-guided behavior (Atiani et al., 2014; Carcea et al., 2017; Elgueda et al., 2019; Francis et al., 2022; Francis et al., 2018; Kuchibhotla et al., 2017), while task experience and perceptual training induce cortical plasticity (Bieszczad & Weinberger, 2010; Caras & Sanes, 2017; Irvine, 2018; Pienkowski & Eggermont, 2011; Polley et al., 2006; Recanzone et al., 1993). Optogenetic excitation or suppression of auditory cortex can bias or impair sound-guided decisions (Ceballo, Piwkowska, et al., 2019; O’Sullivan et al., 2019; Znamenskiy & Zador, 2013) and perceptual learning (Bajo et al., 2019). Importantly, mice can also learn to discriminate imposed cortical activity patterns (Ceballo, Bourg, et al., 2019; Ceballo, Piwkowska, et al., 2019). Directly stimulating auditory cortex isolates the behavioral readout of cortical activity from the subcortical transformations that normally shape cortical responses to sound. It therefore allows a test of whether differences in the rate and strength of optogenetic stimulation can guide behavior. This question is relevant to the broader aim of encoding discriminable information with optical neuroprostheses, including optical cochlear implants (Dieter et al., 2020).

Here, we asked whether mice expressing Channelrhodopsin-2 (ChR2) in excitatory neurons could detect mesoscale photostimulation through a cranial window centered over auditory cortex, distinguish it from external visual and acoustic stimuli, and discriminate differences in photostimulation rate. We further asked whether sensitivity to a weakened optoceptive cue could improve with experience. To investigate, we trained mice in a sequence of head-fixed go/no-go optoception tasks. We found that mesoscale cortical optogenetic stimulation generates a graded internal signal whose behavioral readout depends on stimulation rate, optical power, and experience.

## MATERIALS AND METHODS

### Animals

All animal procedures were performed in accordance with the UMD animal care committee’s regulations. We used 14 mice, aged 13-35 weeks. The behavioral cohort comprised four ChR2-positive males (by01, br01, bb04, wf01) and two ChR2-negative male littermates (mo01, qk01). Optogenetic mice were F1 offspring of CBA/CaJ mice (The Jackson Laboratory, stock 000654) crossed with B6.Cg-Tg(Thy1-COP4/EYFP)18Gfng/J mice (stock 007612). The F1 cross combined ChR2 expression in excitatory neurons with the CBA background, which prevents age-related hearing loss (AHL) associated with the C57BL/6J background (Arenkiel et al., 2007; Fenno et al., 2015; Frisina et al., 2011). The behavioral experiment was not designed to test sex effects since usable mice in our litters were limited to genotypes that were both ChR2-positive and AHL-negative. Thus, only 25% of offspring were expected to qualify, and in our case, all were male.

The extent of auditory cortex covered by our implanted LED (see below) was estimated by reanalyzing widefield calcium imaging data from eight F1 CBA/CaJ × Thy1-GCaMP6s mice (hp01, hs01, op01, db01, to01, je01, sa01, and jz01) from a previously published study (Putnam et al., 2026). The Thy1-GCaMP6s line was C57BL/6J-Tg(Thy1-GCaMP6s)GP4.3Dkim/J (The Jackson Laboratory, stock 024275). The imaging mice were distinct from the six behavioral mice. All mice were housed under a reversed 12 h light/12 h dark cycle.

### Cranial window and LED implantation

Mice received dexamethasone (5 mg/kg, i.p.) at least 1 h before surgery. Anesthesia was induced with 5% isoflurane and maintained at 0.5-2%. Cefazolin (500 mg/kg, s.c.) was administered perioperatively, and body temperature was maintained at 37.5°C with feedback heating. After removal of scalp and connective tissue over the temporal bone, a thin layer of cyanoacrylate (Vetbond, 3M) was applied to the skull and a 3D-printed stainless-steel headplate was fixed at the midline. A 3 mm circular craniotomy was made over auditory cortex, dorsal to the zygomatic arch and anterior to the lambdoid suture near the squamosal-parietal junction. A glass window was placed in the craniotomy, and the space between the glass and brain was sealed with optically transparent silicone elastomer (Kwik-Sil, World Precision Instruments).

For optogenetic photostimulation experiments, a 3 mm diameter, 459 nm blue LED (Chanzon) was fixed directly above the cranial window with optical adhesive (Norland Optical Adhesive). The LED was covered with opaque black heat-shrink tubing so that light was emitted only through the 3 mm face affixed to the cranial window, rather than through the sides of the LED. C&B Metabond dental cement (Parkell) secured the implanted-LED to the skull, which was then painted with black dental cement made by mixing methyl methacrylate with iron oxide powder, further limiting light leakage (Figure 1A). Meloxicam (0.5 mg/kg, s.c.) was given after surgery.

**Figure 1.**
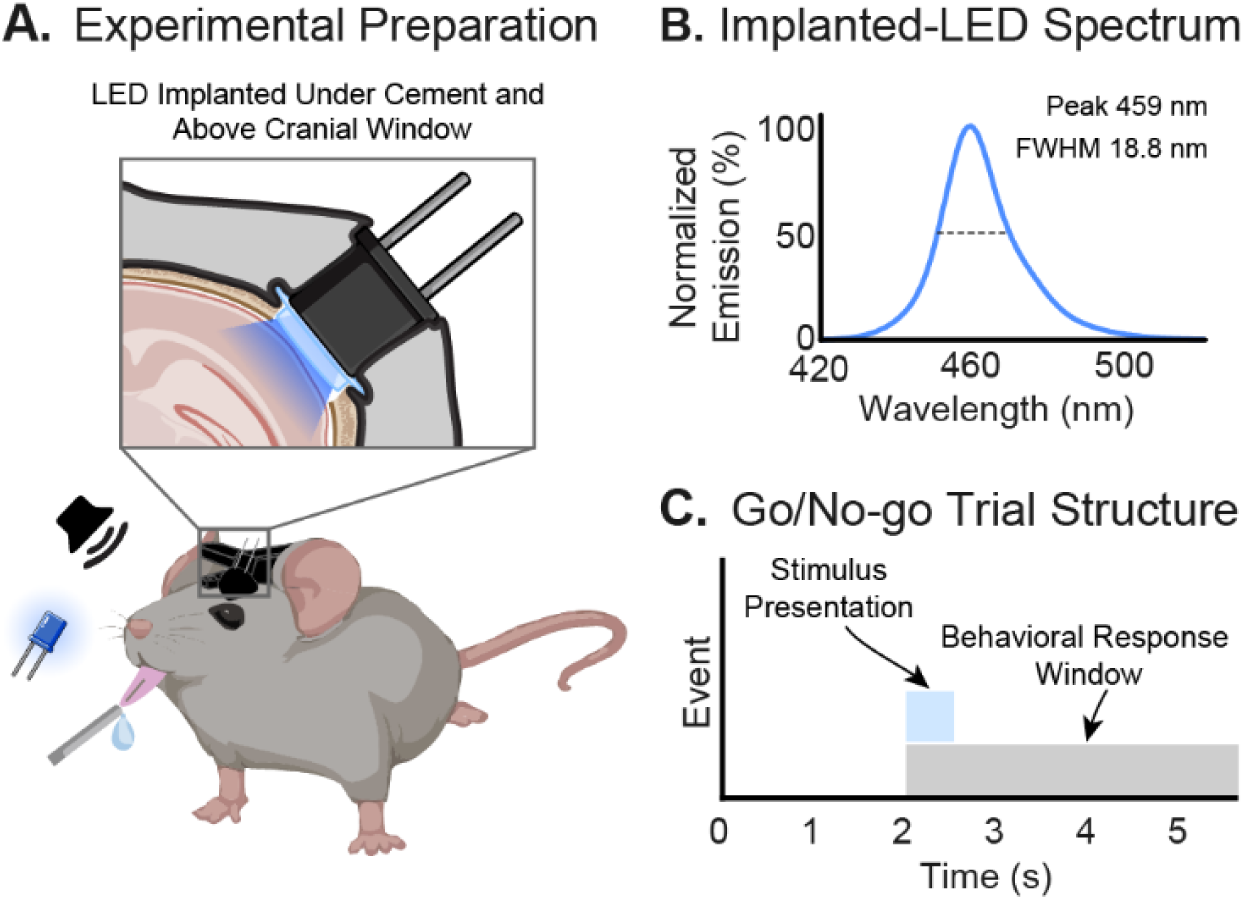
Experimental preparation, LED characterization, and trial structure. **A.** A 3 mm, 459 nm LED was fixed directly above the cranial window and shielded with opaque tubing and black dental cement. Head-restrained mice reported stimulation by licking a waterspout. A separate 459 nm external LED and speaker presented visual and auditory non-targets. **B.** Emission spectrum of the implanted-LED measured at the cranial-window face. The horizontal segment marks the 18.8 nm full width at half maximum (FWHM). **C.** Each trial contained a 2 s prestimulus period, a 0.5 s stimulus interval, and a 3 s poststimulus period. The standard response window extended from stimulus onset through the 5.5 s trial end. A response-window lick on a target trial delivered water; licking on a non-target or LED-off catch trial triggered a 10 s timeout.

The optical characteristics of the LED used both for optogenetic stimulation and as an external visual cue were measured at the LED face, i.e., where it interfaced with the cranial window. The LED emission spectrum was recorded with an ASEQ LR1-B spectrometer (Figure 1B). LED power was measured with a Thorlabs S121C silicon photodiode power sensor. The mean spectrum peaked at 459 nm and had a full width at half maximum of 18.8 nm. The 3-mm-diameter circular window had an area of 7.07 mm². Radiant power for a 5 V drive was 2.55 mW and 1.33 mW for a 2.5 V drive, thus corresponding to incident irradiances of 0.361 and 0.188 mW/mm², respectively.

### Widefield calcium imaging

Widefield GCaMP6s fluorescence was imaged from auditory cortex in awake, head-restrained Thy1-GCaMP6s × CBA mice through the 3 mm cranial window described above. A 473 nm LED provided excitation light, which was spectrally restricted by a 470 nm excitation filter before illuminating the cortex. Emitted fluorescence was collected through a 4× objective, passed through a 505 nm long-pass filter and a 531 nm band-pass filter, and recorded at 5 Hz with 512 × 512-pixel resolution using a CMOS camera controlled by ThorCam software (Thorlabs). After a reference image of the cortical surface was acquired, the focal plane was advanced approximately 750 μm below the surface for functional imaging.

Sounds were presented from an ES1 free-field speaker through an ED1 amplifier (Tucker-Davis Technologies). Neural activity was measured in response to pure-tone frequencies from 2 to 45 kHz at two tones per octave, as well as broadband (4-45 kHz) pulse trains, harmonic complex tones, amplitude-modulated noise, and iterated-ripple noise, with fundamental frequencies (F0s) from 20 to 1280 Hz at two F0s per octave. Each trial contained a 1 s baseline, a 0.5 s sound, and a 2 s post-stimulus period. Each stimulus was repeated 20 times. Interstimulus intervals were 6, 7, or 8 s.

Videos were rigidly registered to the surface reference, spatially filtered, downsampled fourfold, half-wave rectified, and masked to suppress low-signal regions, yielding 128 × 128-pixel maps spanning the window. For each pixel and condition, trials were aligned to sound onset and averaged. ΔF/F was calculated relative to the prestimulus baseline through 1 s after onset. For the present reanalysis, a pixel from a given experiment was classified as auditory responsive if it was assigned a best-frequency or best-F0 for at least one acoustic stimulus class. Masks were combined across stimulus classes and sessions to produce one mask for each mouse. The across-mouse probability map in Figure 2B gives the percentage of mice with an auditory-responsive pixel at each cortical location.

**Figure 2.**
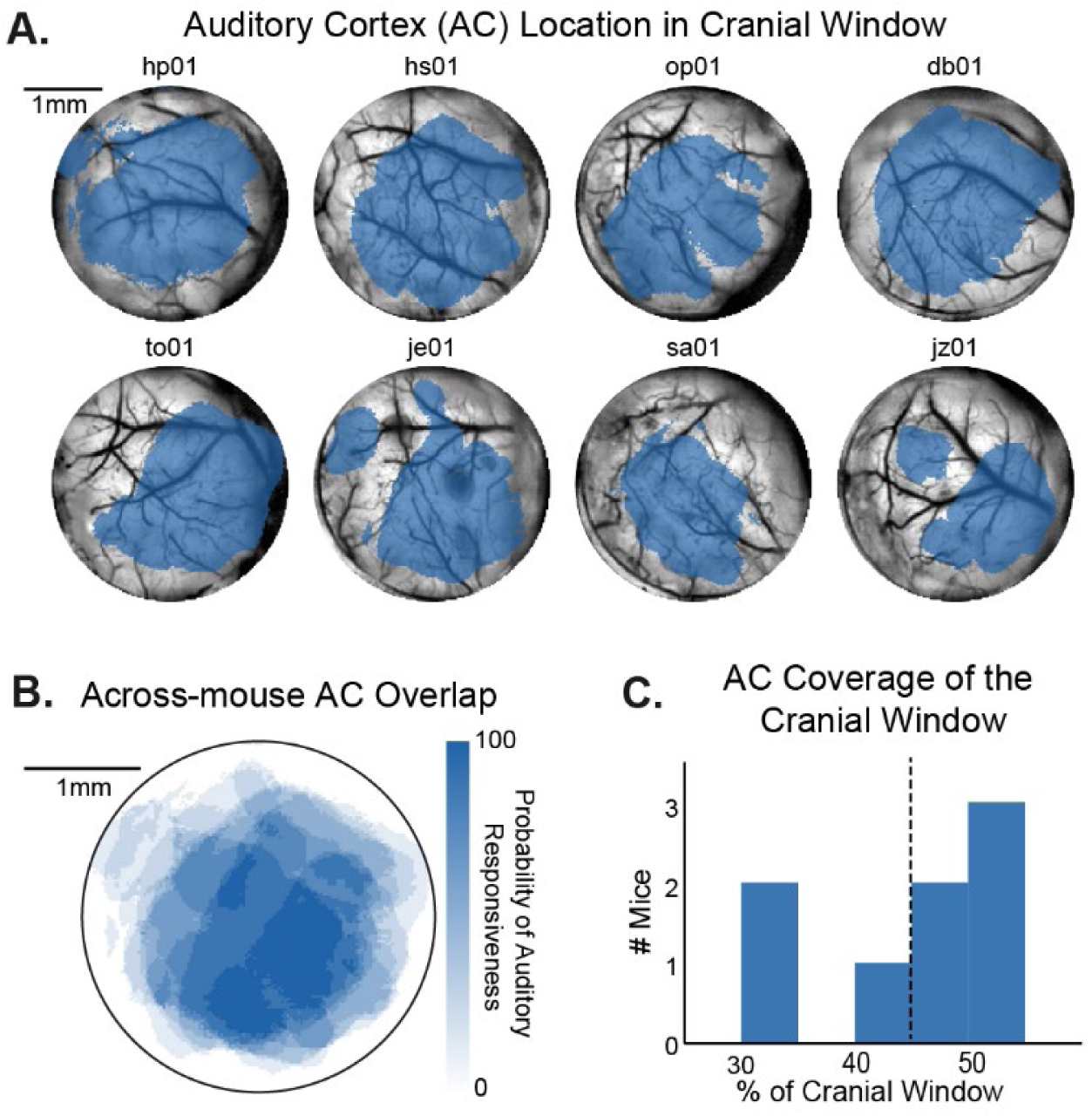
Auditory-responsive cortex within the implanted-LED field. Auditory-responsive coverage was estimated in a cohort of eight Thy1-GCaMP6s × CBA mice. **A.** The top two rows show each mouse’s auditory cortex within the 3 mm cranial window. **B**. Across-mouse auditory cortex overlap: the probability map gives the percentage of mice that had an auditory-responsive pixel at each location. **C.** Auditory cortex coverage of the cranial window: the histogram shows auditory cortical area as a percentage of the circular window; the dashed line marks the across-mouse mean of 44.9%.

### Behavioral apparatus and stimuli

Head-restrained mice performed a sequence of go/no-go tasks while licking a waterspout. A capacitive touch sensor recorded lick times, and custom MATLAB software coordinated the implanted LED, visual and acoustic stimuli, water rewards, and timeouts through a National Instruments NI-6211 data-acquisition board. During Phases 1-6, implanted-LED photostimulation consisted of a 0.5 s train of 25 ms pulses. A 20 Hz train therefore delivered approximately 10 pulses and 250 ms of total illumination, whereas a 10 Hz train delivered approximately five pulses and 125 ms. Phase 7 used 5 ms pulses and presented one of 11 rates from 10 to 20 Hz in 1 Hz increments.

For visual-control trials, a separate 459 nm LED was positioned in front of the mouse and pulsed at 20 Hz. For acoustic-control trials, broadband (4-45 kHz) noise was sinusoidally amplitude modulated at 20 Hz with 100% modulation depth and presented at 75 dB sound pressure level. Acoustic stimuli were synthesized in MATLAB, output at a 200 kHz sampling rate through the NI-6211, amplified by a Tucker-Davis Technologies ED1, and delivered through an ES1 speaker. Speaker output was calibrated and equalized *in situ* at the animal’s head position with a Brüel & Kjær Type 4939 microphone, yielding a flat frequency response from 1 to 64 kHz.

Each trial lasted 5.5 s and comprised a 2 s prestimulus period, a 0.5 s stimulus interval from 2.0 to 2.5 s, and a 3 s poststimulus period (Figure 1C). The behavioral response window began at the stimulus-onset time and continued to the end of the trial. Mice were trained to lick after target (go) stimuli and withhold licking after non-target (no-go) stimuli. A response-window lick on a target trial was scored as a hit and triggered 0.25 s water delivery; no lick was a miss. A response-window lick on a non-target trial was a false alarm and triggered a 10 s timeout; no lick was a correct rejection. An LED-off catch trial preserved normal trial timing and the response window but omitted implanted-LED photostimulation; in Phases 3 and 4, LED-off trials also omitted the external visual or acoustic comparison stimulus. Behaviorally, a catch was a no-go trial: mice had to withhold licking. A response-window lick was a false alarm followed by a 10 s timeout, whereas withholding licking was a correct rejection; no water was delivered. Trials containing a lick before the response window were classified as early-response trials. Their treatment in response-rate and d′ calculations is described below.

### Training sequence

Training proceeded in the fixed sequence illustrated in Figure 3, with trial types interleaved within each phase. During Phases 1-5, the LED delivered 2.55 mW of radiant power at the cranial-window face. Phase 1 was target-only shaping: 100% of trials contained a 20 Hz implanted-LED train. A response-window lick triggered the standard 0.25 s water reward. To establish the stimulus-response association, an additional 0.25 s water was automatically delivered on either 33.3% or 66.7% of trials at 0.5 s after the train ended. Phase 2 trials comprised 66.7% 20 Hz implanted-LED targets and 33.3% LED-off catches. Phase 3 comprised approximately 33.3% each of 20 Hz implanted-LED targets, LED-off catches, and 20 Hz external 459 nm LED non-targets. Phase 4 similarly comprised approximately 33.3% each of 20 Hz implanted-LED targets, LED-off catches, and 20 Hz amplitude-modulated broadband-noise non-targets. Phase 5 comprised 50% 20 Hz target and 50% 10 Hz non-target implanted-LED trains. In Phases 1-5, implanted-LED trains lasted 0.5 s and used 25 ms pulses, yielding approximately 10 pulses at 20 Hz and five pulses at 10 Hz. Phase 6 retained the 50% 20 Hz target and 50% 10 Hz non-target contingency while the LED power was reduced to 1.33 mW. In Phase 7, LED power remained at 1.33 mW and pulse width was reduced to 5 ms. 10-14 Hz photostimulation rates were non-targets, for which a response-window lick triggered a 10 s timeout, and 15-20 Hz were targets, for which a response-window lick triggered water delivery. Phase 7 contained no LED-off catch trials.

**Figure 3.**
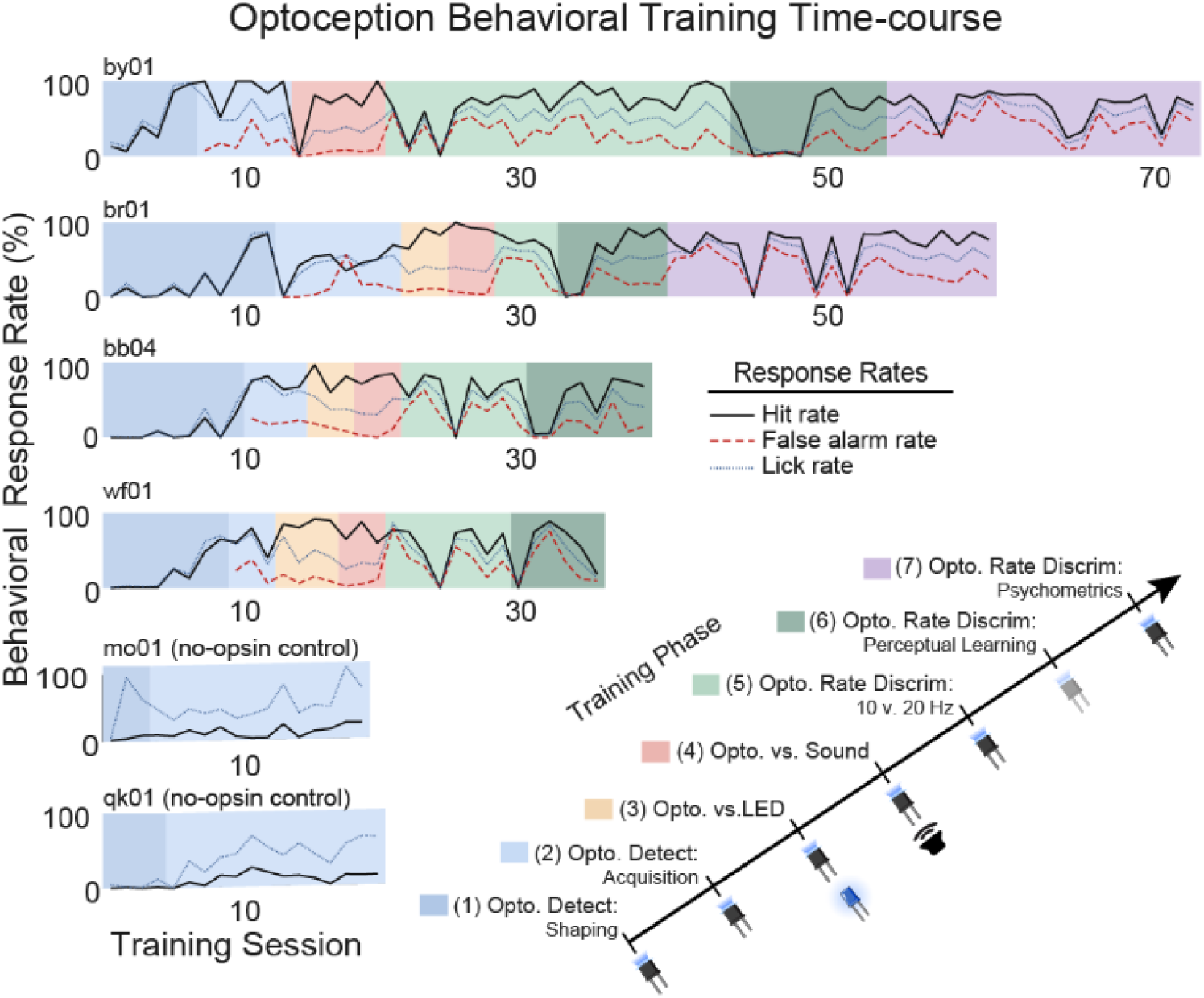
Optoception training across sessions. Session-level hit rate (solid black), false-alarm rate (red dashed), and overall lick rate (blue dotted) are plotted across successive sessions for four ChR2-positive mice (by01, br01, bb04, and wf01) and two ChR2-negative implanted-LED controls (mo01 and qk01). Colored backgrounds show the fixed training sequence. Phase 1 contained only 20 Hz targets. Phase 2 sessions contained 66.7% targets and 33.3% LED-off catches. Phases 3 and 4 contained approximately 33.3% each of targets, catches, and the visual or acoustic non-target. Phases 5 and 6 contained 50% 20 Hz targets and 50% 10 Hz non-targets. In Phase 7, 10-14 Hz were non-targets and 15-20 Hz were targets. The ChR2-negative mice remained in target-only training and did not acquire photostimulation-locked responding.

### Behavioral analysis

Hit rate, false-alarm rate, overall lick rate, and first-lick latency were computed for each session. For ChR2-positive sessions containing both target and non-target trials, hit and false-alarm rates were the fractions of target and non-target trials, respectively, with a response-window lick, after excluding early-response trials. Discrimination sensitivity was calculated as d′ = z(hit rate) - z(false-alarm rate), where z denotes the inverse standard-normal transformation. Before transformation, proportions of 0 and 1 were replaced with 1/(2N) and 1 - 1/(2N), respectively, where N was the denominator of the corresponding response rate. For target-only Phase 1 sessions, the false-alarm term was calculated as the number of early-response trials divided by all recorded trials.

### Statistical analysis

The top row of Figure 4 used one-sided, null-centered nonparametric bootstrap tests of whether the mean session-level sensitivity measure (d′) was below the d′ = 1 training criterion. Each session-level value was expressed as its difference from the criterion, and the observed mean difference was subtracted from every value to create an empirical null distribution centered at zero. Each bootstrap sample consisted of n session-level values drawn with replacement from this null distribution, where n was the number of included sessions. The sample mean was recalculated for each of 20,000 iterations. The one-sided p value was estimated as (b + 1)/(20,000 + 1), where b was the number of null-resampled means less than or equal to the observed mean difference. Mouse-level means and 95% percentile confidence intervals were estimated with 20,000 bootstrap resamples of mice. For the latency distributions in Figure 4, target-trial first licks from 0 to 5.5 s were binned in 50 ms intervals within each mouse and expressed as a percentage of that mouse’s recorded target-trial first licks; lines show across-mouse means and shaded bands show ±2 SEM. Tests were considered significant only when the observed mean was below the criterion and p < 0.05.

**Figure 4.**
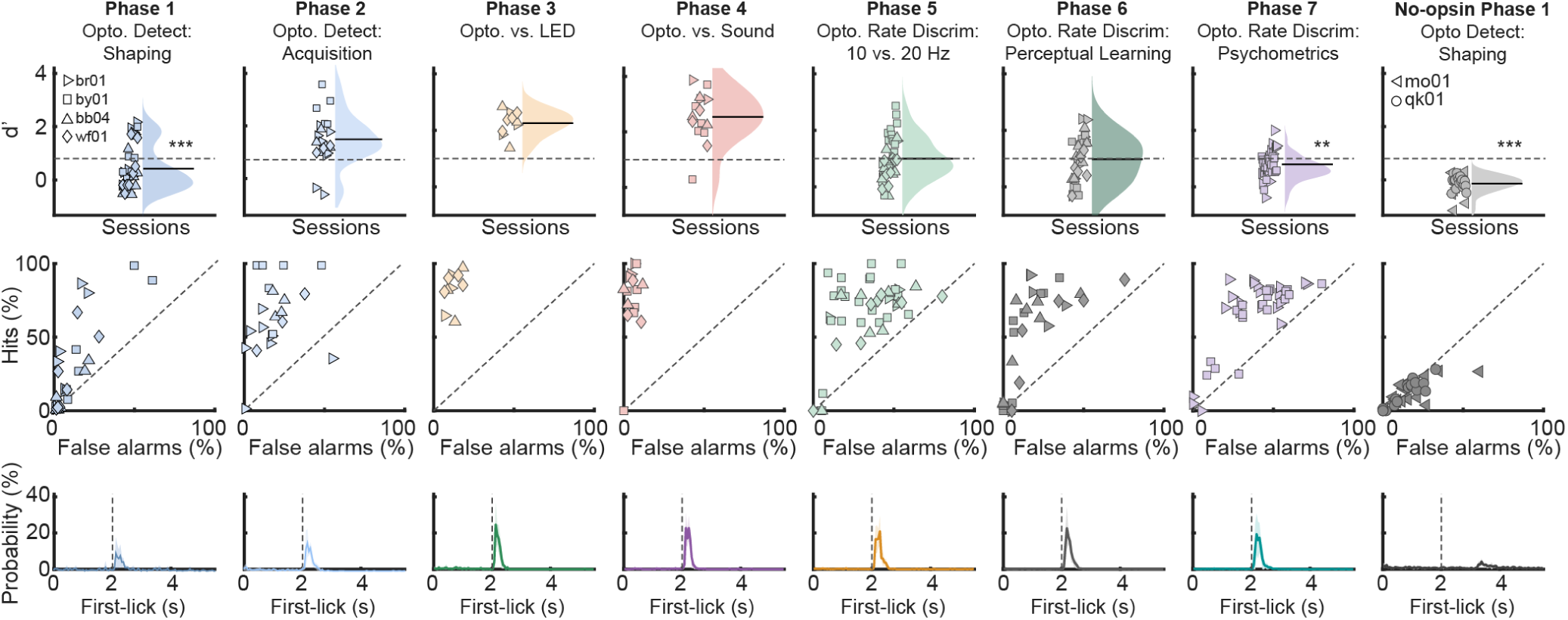
Training Phase Behavior. Columns show individual phases. *Top row,* symbols show individual-session sensitivity (d′). Half-violins show session distributions, and horizontal black lines show panel means. The dashed line marks the d’ criterion of 1. *Middle row,* hit rate versus false-alarm rate for each phase. The diagonal denotes equal rates. *Bottom row,* normalized target-trial first-lick latency distributions. Lines show across-mouse means, shaded bands show ±2 SEM, and the vertical dashed line marks stimulus onset. ChR2-positive n = 4 except external-LED discrimination (n = 3) and psychophysics (n = 2); ChR2-negative n = 2. Symbols show individual behavioral sessions. Asterisks mark panel means significantly below 1 in one-sided, null-centered bootstrap tests with 20,000 resamples (*p < 0.05; **p < 0.01; ***p < 0.001).

To measure the immediate effect of reducing implanted-LED optical power, we used a two-sided paired t test to compare d′ in each mouse’s final higher-power Phase 5 session with its first reduced-power Phase 6 session. To assess perceptual learning after the power reduction, we fit an ordinary least-squares line to session-level d′ as a function of session number for each mouse. The fitted slope estimated the change in d′ per training session, and the conventional two-sided regression test of a zero slope was reported for each mouse in Figure 5A. For the group display in Figure 5B, d′ was averaged across the mice available at each relative Phase 6 session. The plotted group trend was the mean of the animal-specific fitted lines, the shaded envelope shows ±2 SEM across their predicted values, and R² quantifies the agreement between this mean trajectory and the session-aligned group means. Because we predicted that discrimination sensitivity would improve with training, the four mouse-level slopes were tested against zero with a one-sided one-sample t test.

**Figure 5.**
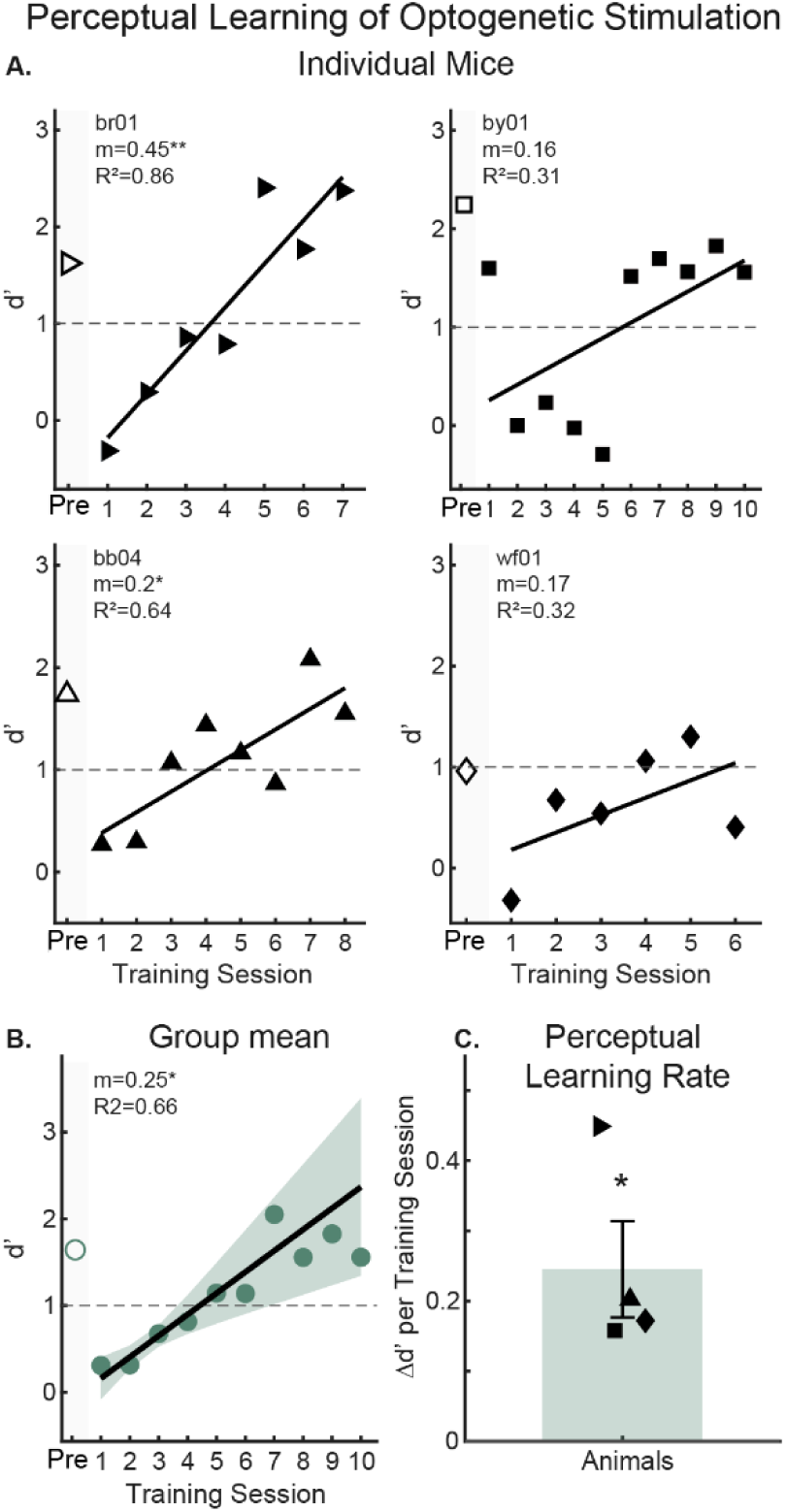
Perceptual learning of a weakened optoceptive cue. **A.** Session d′ for br01, by01, bb04, and wf01. Open symbols at Pre show the final Phase 5 session at the higher LED power, which produced 2.55 mW of radiant power at the cranial-window face; filled symbols show successive Phase 6 sessions after the LED power was reduced to 1.33 mW. In Phase 6, 50% of trials were 20 Hz targets, for which licking delivered water, and 50% were 10 Hz non-targets, for which licking triggered a 10 s timeout. Solid lines are animal-specific ordinary least-squares fits; m is the change in d′ per successive training session and R² is the variance explained. Asterisks beside individual m values indicate two-sided regression tests of a zero slope (*p < 0.05; **p < 0.01). The dashed line marks d′ = 1. **B.** Points show mean d′ across the mice. The line is the mean of the animal-specific fitted lines, and shading shows ±2 SEM across their predicted values. Four mice contributed sessions 1-6, three contributed session 7, two contributed session 8, and one contributed sessions 9-10. **C.** Per-animal learning slopes and mean ± SEM. The mean slope was 0.245 d′ per training session (p = 0.0187). The asterisk beside the group slope in B and above the mean in C denotes this one-sided test of the four animal slopes against zero.

To estimate the psychometric relationship between photostimulation rate and lick probability, we analyzed trials from the last 10 Phase 7 sessions completed by each mouse. Within each mouse, trials were pooled across these 10 sessions at each of the 11 stimulation rates, and lick probability was calculated as the percentage of trials containing a first lick during the response window. The 11 rate-specific probabilities were fit separately for each mouse with a four-parameter logistic function of log2 photostimulation rate: P(z) = L + (U - L)/{1 + exp[-(z - m)/s]}, where z = log2(rate), L and U were the lower and upper asymptotes, m was constrained to the tested log-rate range, and s was constrained to be positive. Parameters were estimated by minimizing the unweighted sum of squared residuals using MATLAB fminsearch, with maximum iteration and function-evaluation counts of 5,000. Goodness of fit was reported as R² = 1 - SSE/SST. The psychometric threshold was defined as the rate at which fitted lick probability lay halfway between L and U. The same function was also fit separately to each of the 10 sessions from each mouse, and session-level thresholds were summarized as the mean ±2 SEM. Figure 6A shows the pooled psychometric functions and session-level threshold insets; Figure 6B shows the fitted lick-probability curves for each of the 10 analyzed sessions from each mouse. All analyses were performed in MATLAB.

**Figure 6.**
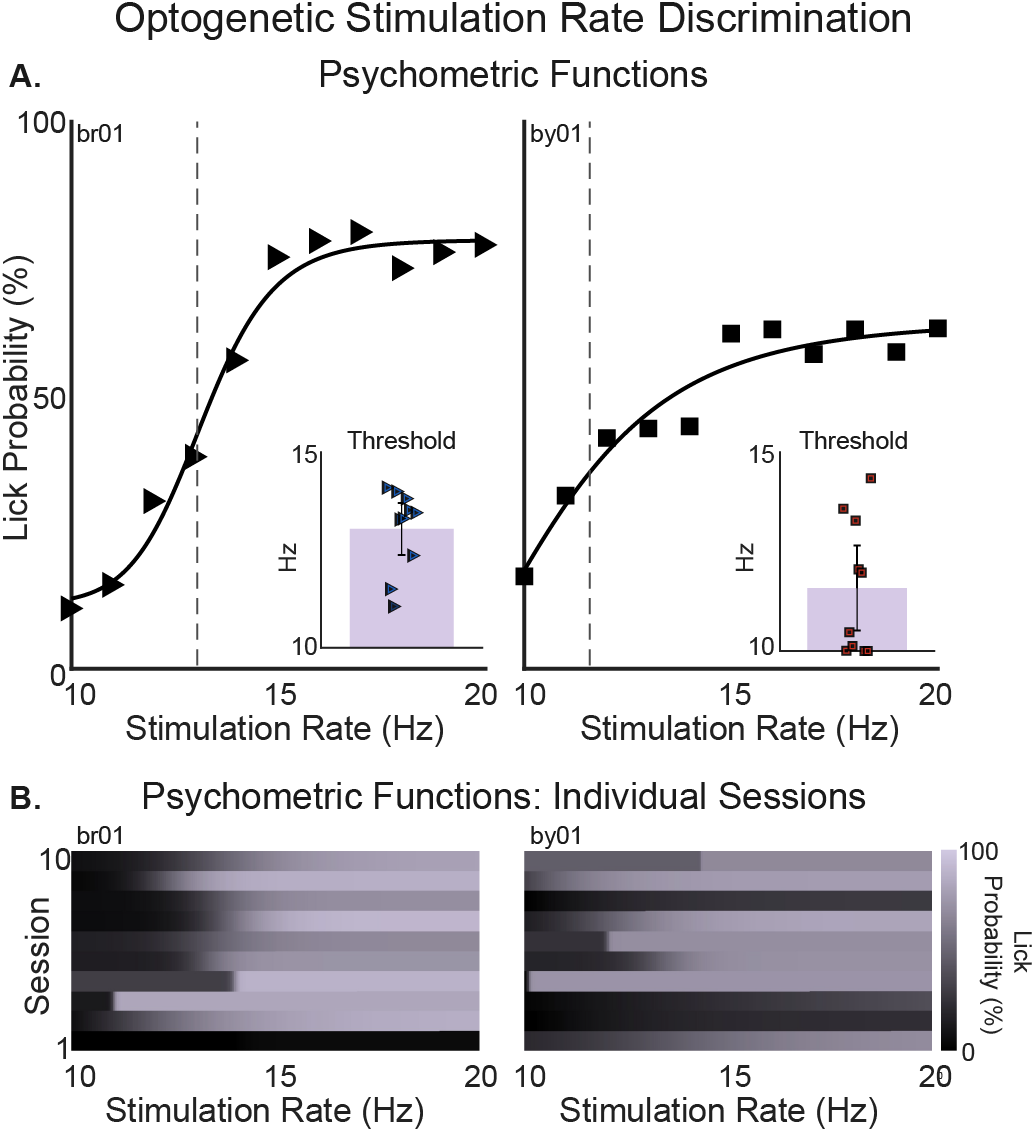
Rate-dependent psychometric functions for optoception. In Phase 7, 10-14 Hz were non-targets and 15-20 Hz were targets. **A.** Symbols show lick probability at each rate after pooling trials across these sessions. Solid curves show least-squares log-rate logistic fits to the pooled values; fitted midpoints were 13.15 Hz for br01 and 10.05 Hz for by01, with R² values of 0.983 and 0.935, respectively. Vertical dashed lines show the mean session-level threshold (13.04 Hz for br01 and 11.58 Hz for by01), defined as the rate halfway between the fitted lower and upper asymptotes. Insets show thresholds from individual session-level fits; bars are means and error bars are ±2 SEM. **B.** Heat maps show fitted lick probability by rate for each of the 10 analyzed sessions from each mouse.

### Data Accessibility

Source data and code for figures ([TBD]) are available in online repositories.

## RESULTS

### Mesoscale photostimulation encompassed auditory cortex

Our experiments aimed to clarify whether ChR2-positive mice could detect mesoscale photostimulation of excitatory neurons through a cranial window centered over auditory cortex. We used an implanted LED as the cortical photostimulation source. The implanted LED emitted a blue spectrum that peaked at 459 nm and had a full width at half maximum of 18.8 nm (Figure 1B). The 3-mm-diameter circular window had an area of 7.07 mm². Radiant power for a 5 V drive to the LED was 2.55 mW and 1.33 mW for a 2.5 V drive, corresponding to incident irradiances of 0.361 and 0.188 mW/mm², respectively.

The 3 mm emitting face of the implanted LED matched the diameter of the cranial window positioned over auditory cortex (Figure 1A), preventing functional imaging of cortex in the behavioral cohort. So, we estimated the distribution of auditory-responsive cortex within our behavioral cohort by analyzing widefield calcium imaging maps of auditory cortex in eight Thy1-GCaMP6s × CBA mice whose 3 mm windows were positioned using the same skull landmarks used here (Figure 2A). Auditory cortex typically occupied 30.5-54.5% of the cranial window (Figure 2C). Across-mouse overlap was greatest in the central and ventral window (Figure 2B). Thus, our implanted-LED preparation enables mesoscale optogenetic perturbation of auditory cortex and nearby regions.

### ChR2-positive mice detected cortical photostimulation and distinguished it from external sensation

To establish whether mesoscale cortical photostimulation generated a behaviorally accessible signal distinct from sensory processing, we asked whether mice could detect photostimulation alone, distinguish it from temporally matched visual and acoustic stimuli, and discriminate differences in photostimulation rate. Training progressed from target-only shaping in Phase 1 to increasingly specific tests in Phases 2-5. In Phase 1, all four ChR2-positive mice received only 20 Hz implanted-LED target trials. Response-window licks were rewarded. To shape the stimulus-response association, water delivery automatically occurred on either 33.3% or 66.7% of trials 0.5 s after the train ended. Phase 1 therefore measured acquisition of stimulus-locked responding. Phase 2 discontinued shaping water and introduced LED-off non-target catch trials. Here, a catch was a no-go trial but without LED photostimulation or an external sensory stimulus. Mouse-level d′ values calculated against the pooled non-target trials were 1.00, 1.60, 1.44, and 1.00 (mean, 1.26; 95% bootstrap CI, 1.00-1.52; Figure 4, second column). Hit rates ranged from 48.7% to 78.2%, whereas false-alarm rates ranged from 15.1% to 20.7%. Thus, all four mice responded more often to 20 Hz cortical targets than to the non-targets.

In Phase 3, three ChR2-positive mice completed sessions in which 20 Hz implanted-LED photostimulation was the target and both LED-off catch trials and a temporally matched 20 Hz external 459 nm LED were non-targets. Implanted-LED targets, catches, and external-LED non-targets accounted for 35.3%, 31.2%, and 33.5%, respectively. Target response rates ranged from 78.1% to 86.8%, compared with 10.0-11.8% on external-LED trials and 8.6-14.6% on LED-off catches. Mouse-level d′ values were 2.09, 1.92, and 2.36 (mean, 2.12; 95% bootstrap CI, 1.92-2.36; Figure 4, third column). The similarly low response rates on catch and external-LED trials show that each mouse distinguished cortical photostimulation from the wavelength- and rate-matched visual stimulus.

In Phase 4, all four ChR2-positive mice completed sessions in which 20 Hz implanted-LED photostimulation was the target and both LED-off catches and 20 Hz amplitude-modulated broadband noise were non-targets. Implanted-LED targets, catches, and acoustic non-targets accounted for 33.8%, 32.6%, and 33.6% of trials, respectively. Target response rates ranged from 68.8% to 94.8%, compared with 2.7-5.8% on acoustic non-target trials and 2.5-9.3% on LED-off catches. Mouse-level d′ values were 3.29, 2.25, 2.56, and 2.02 (mean, 2.53; 95% bootstrap CI, 2.13-3.03; Figure 4, fourth column). Thus, every mouse responded selectively to cortical photostimulation over both the temporally matched acoustic stimulus and no-stimulation catches.

Phase 5 tested whether mice were sensitive to the pulse rate of cortical photostimulation. 20 Hz implanted-LED targets and 10 Hz implanted-LED non-targets each accounted for 50% of trials. Mouse-level d′ values were 0.88, 1.15, 0.62, and 0.63 (mean, 0.82; 95% bootstrap CI, 0.62-1.02; Figure 4, fifth column). While only one mouse reached d′ > 1, every mouse had a positive d′, consistent with discrimination between the two stimulation rates.

Finally, we asked whether ChR2-negative mice could detect photostimulation by the implanted LED, as a control for non-opsin-mediated effects in ChR2-positive mice. Two ChR2-negative littermates failed to acquire stimulus-locked responding. Their d′ values were -0.123 and -0.118, and response-window lick rates were 13.2% and 14.8% (Figure 4, eighth column). Across the ChR2-positive conditions, first-lick distributions increased sharply after stimulus onset, whereas the control distributions lacked a comparable stimulus-locked peak; instead, a small peak occurred only when shaping water was automatically delivered (Figure 4, bottom row). These control results establish that opsin-mediated cortical activation carried the behaviorally meaningful cue in ChR2-positive mice, rather than presentation of the implanted LED alone.

### Perceptual learning improved optoceptive sensitivity

Perceptual learning predicts that experience can improve the behavioral readout of an optoceptive cue after that cue is weakened. Phase 6 tested this prediction by reducing the implanted-LED power from 2.55 to 1.33 mW at the cranial-window face while retaining the learned 20 Hz target versus 10 Hz non-target contingency. 50.2% were 20 Hz targets and 49.8% were 10 Hz non-targets. The weaker stimulation initially reduced discrimination sensitivity in every ChR2-positive mouse (Figure 5A): between the final Phase 5 session and the first Phase 6 session, d′ changed from 1.62 to -0.32, 2.25 to 1.60, 1.74 to 0.27, and 0.96 to -0.32. The mean within-mouse change was -1.33 d′ units (t test, p = 0.0155). This immediate decrease established that the reduced-power stimulus provided a weaker optoceptive cue.

With continued experience, optoceptive sensitivity increased across Phase 6 sessions. Fitted slopes were 0.45, 0.16, 0.20, and 0.17 d′ per session, all positive; the individual fits explained 31-86% of the variance in session-level sensitivity (Figure 5A). The mean animal-level slope was 0.245 d′ per session (t test, p = 0.0187), and the corresponding mean fitted trajectory explained 66% of the variance in the session-aligned group means (Figure 5B,C). Because both optical power and the task contingency remained fixed throughout Phase 6, increased sensitivity to the weakened optoceptive cue demonstrates perceptual learning across sessions.

### Optoception showed a psychometric dependence on photostimulation rate

If photostimulation rate is available as a graded optoceptive cue, intermediate rates should systematically bias behavioral choices. Phase 7 tested this prediction with a denser set of photostimulation rates. 11 integer rates from 10 to 20 Hz were randomly interleaved; 10-14 Hz non-target rates accounted for 45.6% of trials and the 15-20 Hz target rates for 54.4%. Lick probability showed a saturating rate dependence in both mice, increasing primarily from 10 to 15 Hz and remaining near an upper plateau from 15 to 20 Hz (Figure 6). Logistic fits to the pooled rate-specific lick probabilities accounted for 98.3% and 93.5% of the variance for br01 and by01, respectively (Figure 6A). Separate fits to each session yielded mean thresholds of 13.04 ± 0.66 Hz for br01 and 11.58 ± 1.07 Hz for by01 (mean ±2 SEM across 10 sessions per mouse; Figure 6A, insets). Thus, cortical photostimulation rate acted as a graded perceptual cue.

## DISCUSSION

Mesoscale cortical photostimulation generated a behaviorally meaningful internal signal: mice distinguished it from matched external sensations, behavioral responses varied with photostimulation rate and optical power, and mice learned to use the signal more effectively with experience. ChR2-positive mice detected 20 Hz stimulation against LED-off catches, distinguished photostimulation from temporally matched visual and acoustic controls, and discriminated 20 from 10 Hz trains. Reducing optical power immediately decreased sensitivity, whereas continued training at the reduced power increased sensitivity across sessions. Together, these results establish optoception as a psychophysically tractable consequence of mesoscale cortical activation.

The rate dependence observed here extends prior work on detection of sparse cortical activation (Dalgleish et al., 2020; Huber et al., 2008), discrimination of photostimulation pulse count (Pancholi et al., 2023), and temporal judgments based on electrical microstimulation (Romo et al., 2000; Romo et al., 1998; Yang et al., 2008). Optogenetic detection thresholds for novel V1 stimulation can improve across sessions (Akitake et al., 2023). Here, Phase 6 tested whether comparable perceptual learning occurs for optoceptive rate discrimination after an abrupt reduction in optical power. Lowering optical power immediately reduced sensitivity in every mouse, establishing that the cue had weakened. Sensitivity then increased while optical power and the 10-versus-20 Hz contingency remained unchanged, showing improvement in how the weakened artificial cortical signal was used for choice. Auditory cortex also contributes causally to perceptual learning for external acoustic cues (Bajo et al., 2019; Caras & Sanes, 2017), whereas the present result shows learning of an artificial cue generated by direct cortical activation. The improvement could reflect more efficient downstream readout, plasticity within the stimulated network, or both.

Our preparation enables trial-resolved optoception measurements with cost-efficient hardware. A commercial LED was bonded to a chronic cranial window and shielded with opaque tubing and dental cement. Compared with patterned optical stimulation systems (Carrillo-Reid et al., 2019; Chong et al., 2020; Gill et al., 2020; Kang et al., 2025; Marshel et al., 2019) or fabricated multisite cortical μLED arrays (Greer et al., 2026), the single-emitter design requires neither beam alignment nor a custom patterned array. Its broad illumination is well suited to testing whether a mesoscale perturbation is reportable and how that report changes with stimulation parameters and training. The same geometry is wavelength-extensible. An emitter matched to red-shifted excitatory opsins such as ReaChR or Chrimson could increase spectral flexibility and tissue penetration (Klapoetke et al., 2014; Lin et al., 2013), whereas inhibitory opsins such as Jaws or ArchT could extend the assay to optical suppression (Chuong et al., 2014; Han et al., 2011). In summary, mesoscale cortical optogenetic stimulation generated a graded artificial perceptual cue that mice distinguished from matched sensory events and learned to use more effectively after optical power was reduced.

## AUTHOR CONTRIBUTIONS

G.S. and N.A.F. designed and performed research; N.A.F. analyzed data. G.S. and N.A.F. interpreted results and wrote the manuscript.

## ACKNOWLEDGMENTS

This work was supported by a University of Maryland Brain and Behavior Institute seed grant to N.A.F.

